# Protocol for open-source Automated Universal high-content multiplex fluorescence for RNA *in situ* Analysis (AURA)

**DOI:** 10.1101/2024.06.28.601140

**Authors:** Jean Descarpentrie, Florian Bernard, Wilfried Souleyreau, Lucie Brisson, Thomas Mathivet, Ioannis S. Pateras, Océane C. B. Martin, Maria Lopez Chiloexhes, Teresa Frisan

## Abstract

In situ hybridization visualizes RNA in cells, but image analysis is complex. We present a protocol based on open-source software for automated high-content multiplex fluorescence *in situ* transcriptomics analysis. Steps include nuclei segmentation with a Fiji macro and quantification of up to 14 mRNA probes per image. We describe procedures for storing raw data, quality control images and the use of a Python app to summarize all the results in one spreadsheet detailing the number of single or co-positive cells.

## Before you begin

### Perform *in situ* transcriptomic analysis and imaging

**Timing: Variable, depending on the method of choice for *in situ* transcriptomic analysis**

1. Perform the *in situ* transcriptomic analysis using the method of choice. **Note:** The protocol has been tested for the RNAscope multiplex V2 assay, but it applies to any equivalent analysis involving intracellular fluorescent and chromogenic specks-like staining, extending its utility beyond *in situ* transcriptomics, and appealing to a wide scientific community.
2. Acquire images by confocal microscopy or tissue slide scanner. **Note**: We recommend acquiring between 700 and 1000 high-quality nuclei per sample. **Note**: Image analysis can also be performed on images acquired with a tissue slide scanner for whole slide imaging, as long as the noise-to-signal ratio is low, and the nuclei and dots are clearly detectable by the experimenter. As a rule of thumb, nuclei and dots should be clearly visible in the image. We do not recommend using epifluorescence microscopy due to its high noise-to-signal ratio. A magnification of 40x is recommended for digitization of *in situ* hybridization ^1^. For confocal imaging, 63x magnification can be used, but this will require acquiring a larger number of images per sample to ensure a statistically representative set. Tissue slide scanners may allow further image magnification using the scanner software.
3. Download the Fiji software at https://imagej.net/software/fiji/#downloads within a Linux, MacOS, or Windows operative system.
4. If your acquired images are not in .TIFF format, convert them using Fiji.

### Install plugins and macro in Fiji

**Timing: 5 minutes**

**Note:** We have developed two macros: **AURA** and **AURA light**.

AURA performs a batch analysis, resulting in a spreadsheet file containing the quantification results for all images in the selected folder. AURA light helps users to identify quickly the parameters for the correct nuclear and dot segmentation for the analyzed tissue on a selected image. This allows for rapid parameters’ adjustment in case of erroneous segmentation. AURA light does not support batch analysis.

5. Download AURA light and AURA macro and plugins
  a. To download the macros go to https://github.com/FlorianBrnrd/aura-data-processing, open the “AURA light” (AURA_Light_v5.ijm) or “AURA” folder (AURA_macro_v1.5.ijm), click on the download command (Figure 1A and 1B, red rectangle).
  b. To download the plugins go to https://mcib3d.frama.io/3d-suite-imagej/#download, click on the “**Download Bundle”** link, and download the 3D ImageJ Suite.
6. Install the plugins and the macro
  a. Open the Fiji folder: for MacOS use the secondary click on the Fiji application and select the Show Package Contents option (Figure 1C, red circle), for Windows users select the Fiji app folder. Proceed with steps b) and c) for both operative systems. Figure 1D shows the content of the Fiji app folder.
  b. Drag the downloaded AURA light macro (AURA_Light_v5.ijm) or AURA macro (AURA_macro_v1.5.ijm) files (purple rectangle in Figure 1E) into the “toolsets” folder (Figure 1D, purple circle) contained in the “macros folder” (Figure 1D, red circle).
  c. Drag the three plugins, marked with a blue rectangle, from the downloaded folder “3D ImageJ Suite” (Figure 1F) into the “plugins” folder (Figure 1D, blue circle).

**Figure 1.**
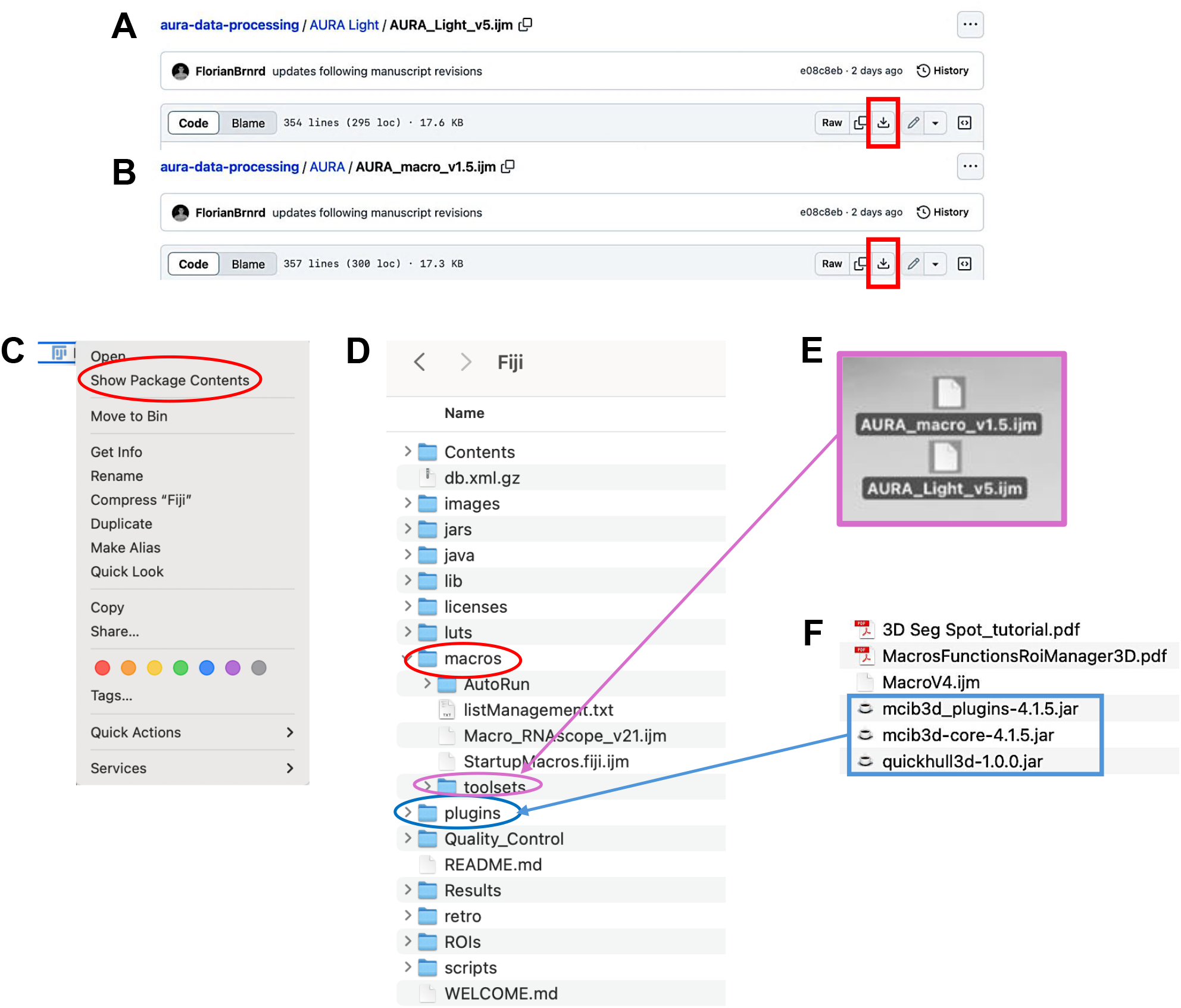
Install the AURA light and AURA macro and plugins. Screenshot of the AURA light **(A)** and AURA macro code **(B)**, the red rectangle indicates the download command. (**C**) Screenshot of the secondary click on the Fiji app for MacOs. (**D**) Screenshot of the Fiji folder content. **(E)** the AURA light and AURA macro files (marked with a purple rectangle) are dragged into the “toolset” folder (purple circle) contained within the “macros” folder (red circle). **(F)** the Fiji Plugins, contained in the downloaded folder “3D ImageJ Suite” (blue rectangle), are dragged into the “plugins” folder (blue circle).

### Key resources table

| REAGENT or RESOURCE | SOURCE | IDENTIFIER |
| --- | --- | --- |
| **Software and algorithms** |  |  |
| ImageJ/Fiji | Schindelin et al.<sup>2</sup>, Schneider et al.<sup>3</sup> | https://imagej.net/software/fiji/#downloads |
| 3D ImageJ Suite Plugins | Ollion et al.<sup>4</sup> | https://mcib3d.frama.io/3d-suite- |
| AURA and AURA light macros | This work | https://github.com/FlorianBrnrd/a |
| LibreOffice | The Document Foundation | https://www.libreoffice.org/ |
| **Other** |  |  |
| Sample slides processed for *in situ* fluorescence transcriptomic | Not required | According to the users protocol |
| Confocal Microscope or Tissue slide scanner to acquire the sample images | Not required | According to the users protocol |

### Step-by-step method details

This step includes the analysis of a single image to define the segmentation parameters.

**Note:** If the parameters have been predefined by the user, steps 1 to 5 can be skipped, and the user can proceed directly to step 6.

**Timing: maximum 30 min**

1. Make a separate folder with a representative image of your tissue.
2. Launch AURA light to define the setting for segmentation.
  a. Click the red double arrow in the Fiji toolbar (Figure 2A, red rectangle) and select the AURA light macro from the drop-down menu. **Note:** the AURA light macro shortcut will appear in the Fiji menu toolbar (Figure 2B, red rectangle).
  b. Click on the AURA light macro shortcut button (Figure 2B, red rectangle).
3. Select the folder containing the images to be analysed. **Note:** The macro begins by prompting the user to select the folder containing the image to be analyzed. The tool cannot perform batch analysis; therefore, we recommend setting a folder with just one image. **Note:** Images must be in .TIFF format.
4. Input the analysis settings in the macro dialogue box (Figure 3)
  a. Choose the number of channels, including the channel used for nucleus staining, that you want to analyze (minimum 2 channels and maximum 15 channels) (Figure 3A).
  b. Name your channel based on the specific marker that you used (Figure 3B). **Note:** Do not use special characters: % $ ! & / \ : ; « » : % & # @.
  c. Choose the channel used for nucleus staining amongst all channels (Figure 3C).
  d. Choose the maximum expansion area around nuclei (Figure 3D), which is used to identify individual cells and quantify the number of dots within a cell.
  e. Select the channels to be analyzed (Figure 3E). **Note :** This tool allows maximum flexibility in channel analysis. **Note:** The settings will be saved at the end of the analysis in the filename “Analysis_Settings.txt” (Figure 4A).
  f. Based on your negative control image choose the minimum dot size for the analysis for each channel (Figure 3F). **Note:** This procedure is required if the negative control has a few dots smaller than the positive control. These background dots can be used as a reference to choose the size tolerance. The default pre-set value is 0,1μm (Figure 3F). If no background is detected in the negative control the threshold for size dot can be changed to 0, and all dots in the test sample will be segmented and counted.
  g. Click “OK” on the dialogue that will open when the analysis has been completed **Note:** The images with the set segmentation parameters will be available in the same folder chosen in step 3 (Figure 4B). Figure 5 shows two examples of nuclei segmentation using two expansion values. **Note:** If the segmentation is satisfactory, proceed to the next step of the protocol. If not, the macro can be re-run by changing the parameters. For each run, a separate analysis .txt file and QC image will be saved as a record (Figure 4B).

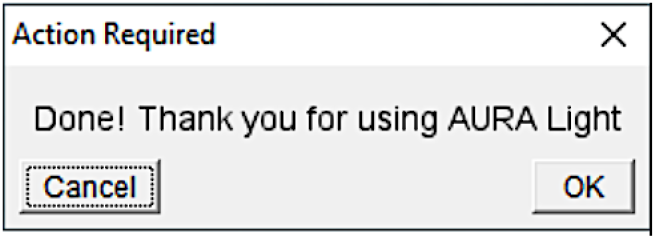

**Figure 2.**
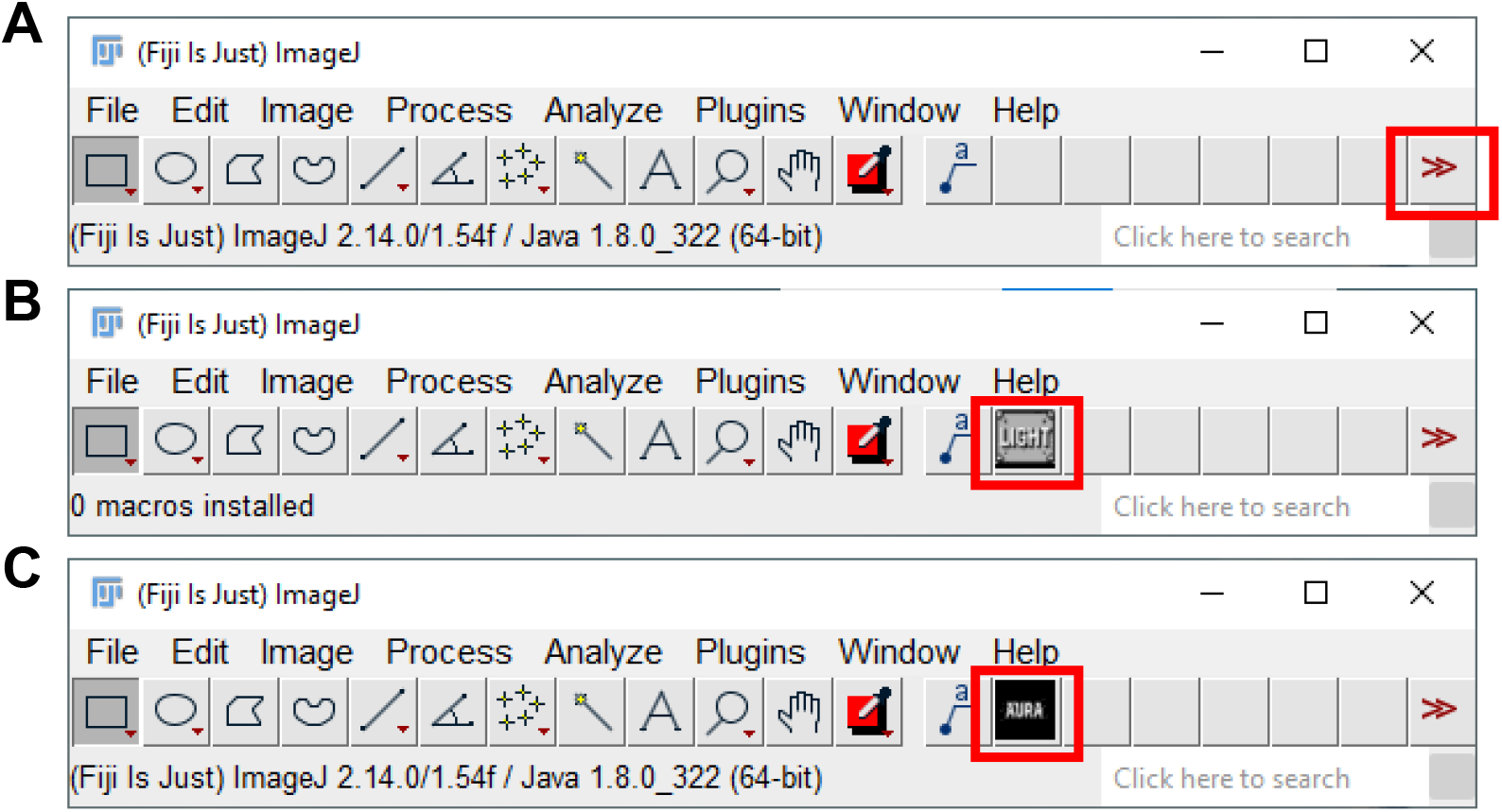
Fiji interface. **(A)** Selection of the AURA macro from the dropdown menu (red rectangle). **(B)** AURA light macro shortcut (red rectangle). (**C**) AURA macro shortcut (red rectangle).

**Figure 3.**
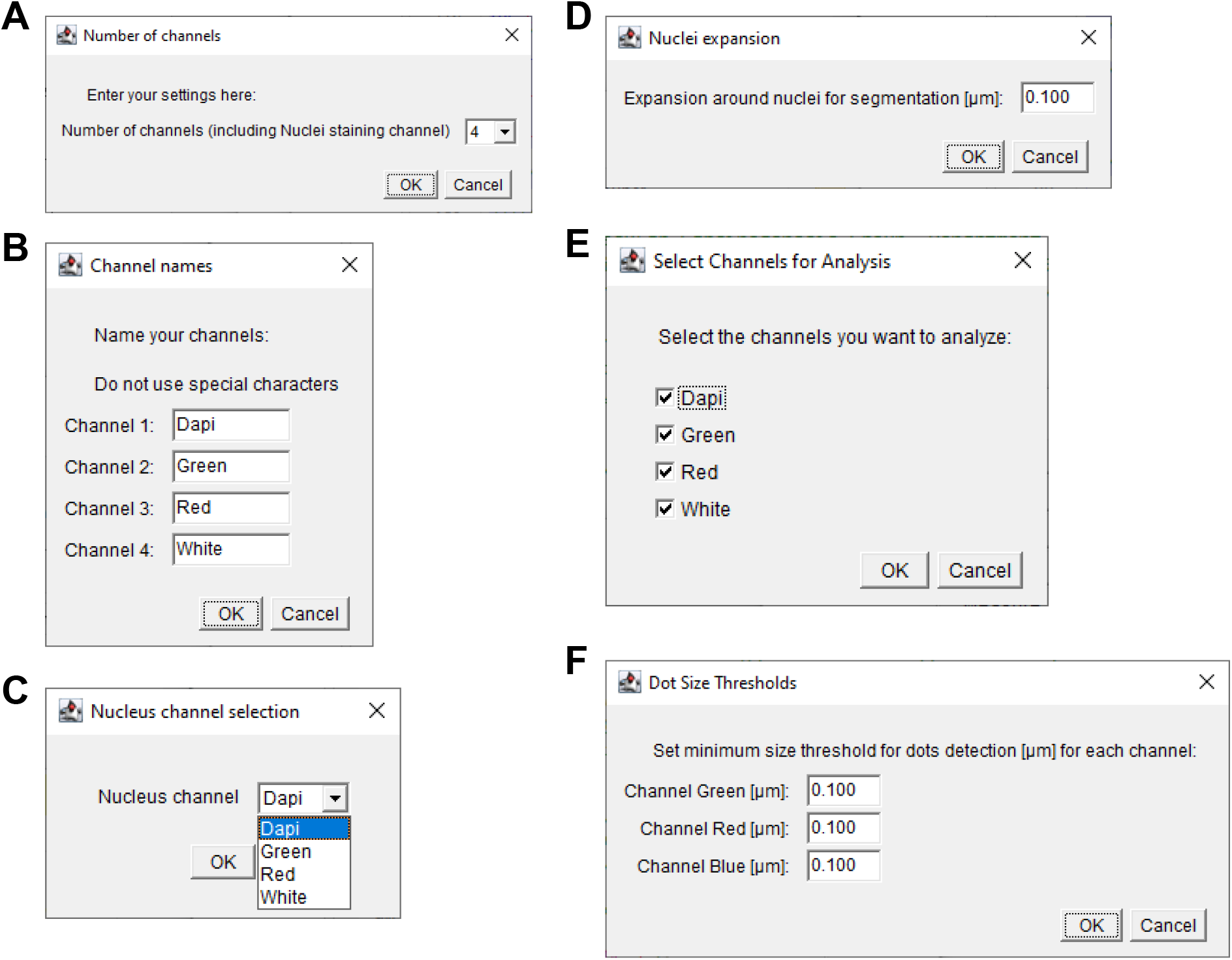
Macro options dialog used to configure settings for image analysis. **(A)** Selection of the number of channels: indicates the total number of channels used in the analysis, including the nuclei staining channel, minimum 2 and maximum 15. **(B)** Channels Options Dialog. Users can label each channel used in the analysis. **(C)** Nucleus Channel Selection Dialog. The user selects the channel used for nuclei identification inside a dropdown menu. **(D)** This dialog box allows users to set parameters for nucleus segmentation. Expansion around nuclei for segmentation [µm]: specify the expansion radius around the nuclei for segmentation, set to 0.100 μm (default setting), and can be changed on the dialogue if required. **(E)** Selection of the channels to be analyzed. **(F)** Definition of the Dot size threshold that can be set for each channel selected in (A). The minimum dot size [μm]: defines the minimum size threshold for the mRNA transcript dots to be considered in the analysis, set to 0.100 μm (default setting), and can be changed on the dialogue if required.

**Figure 4.**
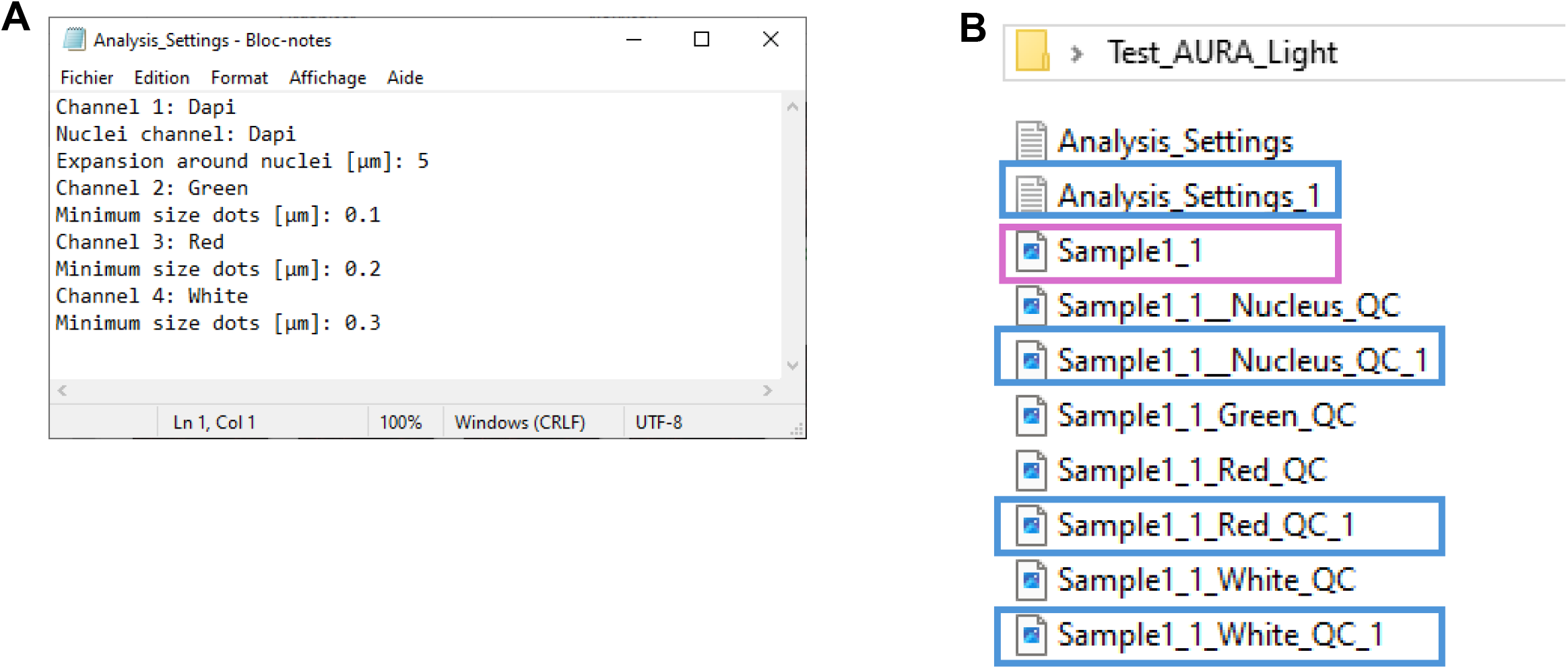
Test folder AURA light. **(A)** The Analysis_Settings .txt file contains all the details for the segmentation parameters. **(B)** The test folder selected for AURA light processing containing: i) the image selected for the analysis (purple rectangle); ii) the two settings tested summarized in .txt files (Analysis _setting and Analysis_Setting_1), and the quality control images with the two settings tested. The blue rectangles indicate the QC files from the second analysis (Analysis_ Setting_1).

**Figure 5.**
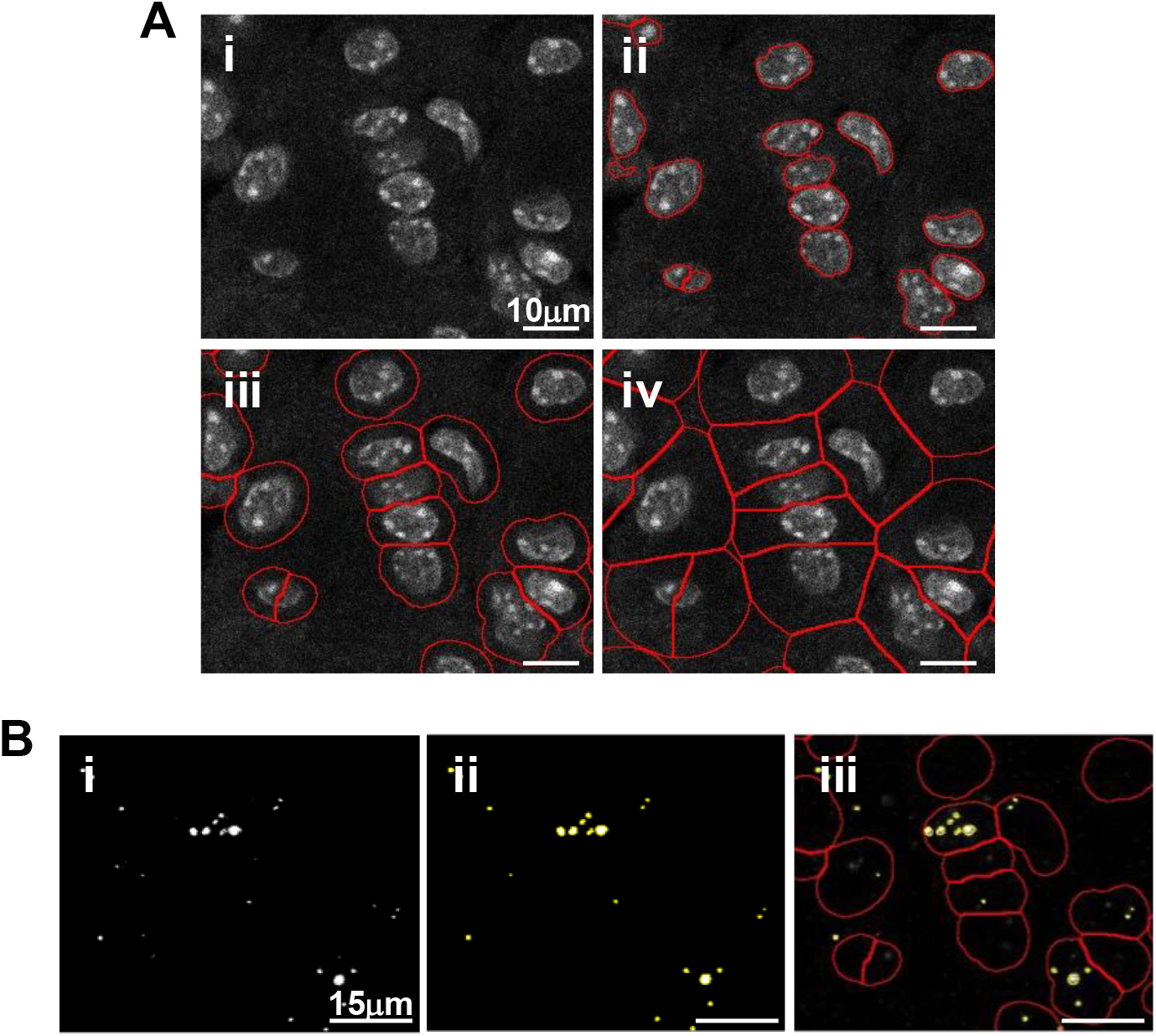
Segmentation results. **(A)** Example of nuclei segmentation process and cell area identification: i) nuclei staining; ii) nuclei segmentation; iii) results of segmentation with an expansion around nuclei of maximum 2 µm; iv) results of segmentation with an expansion around nuclei of maximum 7 µm. **(B)** Example of dots segmentation process and dot analysis: i) Representative micrograph of fluorescent dots; ii) Dots segmentation highlighted in yellow; iii) Dots counting per cell based on the defined cell area.

### Fiji macro analysis with AURA

This step includes the batch analysis of an image set.

**Timing: 1 min per 10 images at magnification 40x (higher magnification will take longer time)**

6. Launch AURA
  a. Click the red double arrow in the Fiji toolbar (Figure 2A, red rectangle) and select the AURA macro from the drop-down menu.
  b. The AURA macro shortcut will appear in the Fiji menu toolbar (Figure 2C, red rectangle).
  c. Click on the AURA macro shortcut button (Figure 2C, red rectangle).
  d. Select the folder containing the images to be analysed **Note:** The macro begins by prompting the user to select the folder containing the images to be analysed. The tools can perform batch analysis, thus all the images in the selected folder will be evaluated. The final .CSV format files (step 9 “Step-by-step method details”) will also be saved in the selected folder. **Note:** Images must be in .TIFF format. The macro itself does not have image size limitations. However, the ability to open and handle large images is determined by Fiji’s capacity (https://imagej.net/learn/faq) and the computational power of the user’s hardware.
7 Input the analysis settings in the macro dialogue box (Figure 3)
  a. Choose the number of channels, including the channel used for nucleus staining that you want to analyze (minimum 2 channels and maximum 15 channels) (Figure 3A).
  b. Name your channel based on the specific marker that you used (Figure 3B). **Note:** Do not use special characters: % $ ! & / \ : ; « » : % & # @.
  c. Choose the channel used for nucleus staining amongst all channels (Figure 3C).
  d. Choose the maximum expansion area around nuclei (Figure 3D), which is used to identify individual cells and quantify the number of dots within a cell (Figure 5A and 5B). **Note:** The default pre-set value is 0,1μm, but the expansion depends on the cell size and tissue type. We recommend testing several expansion values on a few images to find the correct value for the segmentation of your samples using AURA light before starting the full analysis (steps 1 to 5). Staining with surface markers may help define a suitable expansion value. Presently the AURA macro does not allow to curate manually the expansion of individual nuclei in an image.
  e. Select the channel to be analyzed (Figure 3E). **Note:** This tool allows maximum flexibility in channels’ analysis. **Note:** The settings will be saved at the end of the analysis in the filename “Analysis_Settings.txt” (Figure 4A).
  f. Based on your negative control image choose the minimum dot size tolerance for the analysis for each channel (Figure 3F). **Note:** This procedure is required if the negative control has a few dots smaller than the positive control. These background dots can be used as a reference to choose the size tolerance. The default pre-set value is 0,1μm. If no background is detected in the negative control the threshold for size dot can be changed to 0, and all dots in the test sample will be segmented and counted.
  8. Click “OK” on the dialogue that will open when the analysis has been completed
  9. Visualize the macro-output saved in the following three sub-folders, present inside your Images folder (Figure 6):
    a. Quality_Control
    b. Results
    c. Regions of interest (ROIs) **Note:** The Quality control folder contains already processed quality control images for each image (Figure 7A), including:
      a. Nuclei segmentation, marked in red on the nuclei channel image to assess segmentation quality (Figure 7Bi).
      b. Counted dots, marked in yellow and showed in association with the nuclei segmentation (in red) for each channel (Figure 7Bii).

The Results folder (Figure 8) contains:

a. .CSV files generated for each channel per analyzed image (Figure 8A).
b. A single .TXT file that summarizes the analysis settings selected in Step 7 (Figure 8A).

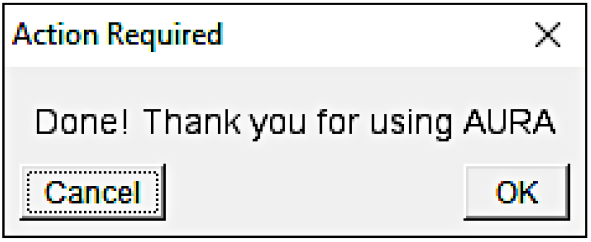

The ROIs folder (Figure 8B) contains the raw data for the regions of interest generated by the macro, including nuclei labelling. To visualize the raw data, first drag the RoiSet file, followed by the corresponding .TIFF image file for the selected channel, into the Fiji interface (Figure 8C). The ROI Manager drop-down menu will appear (Figure 8D, left panel). To display the segmentation, select “Show All” and “Labels”. The image corresponding to the selected ROI will open (Figure 8D, right panel).

**Figure 6.**
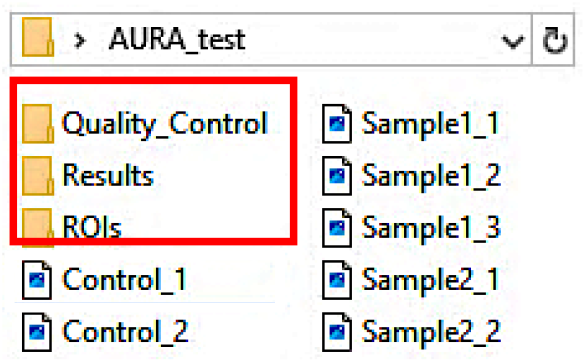
AURA subfolders. Representative image of the three subfolders generated by AURA, marked with a red square.

**Figure 7.**
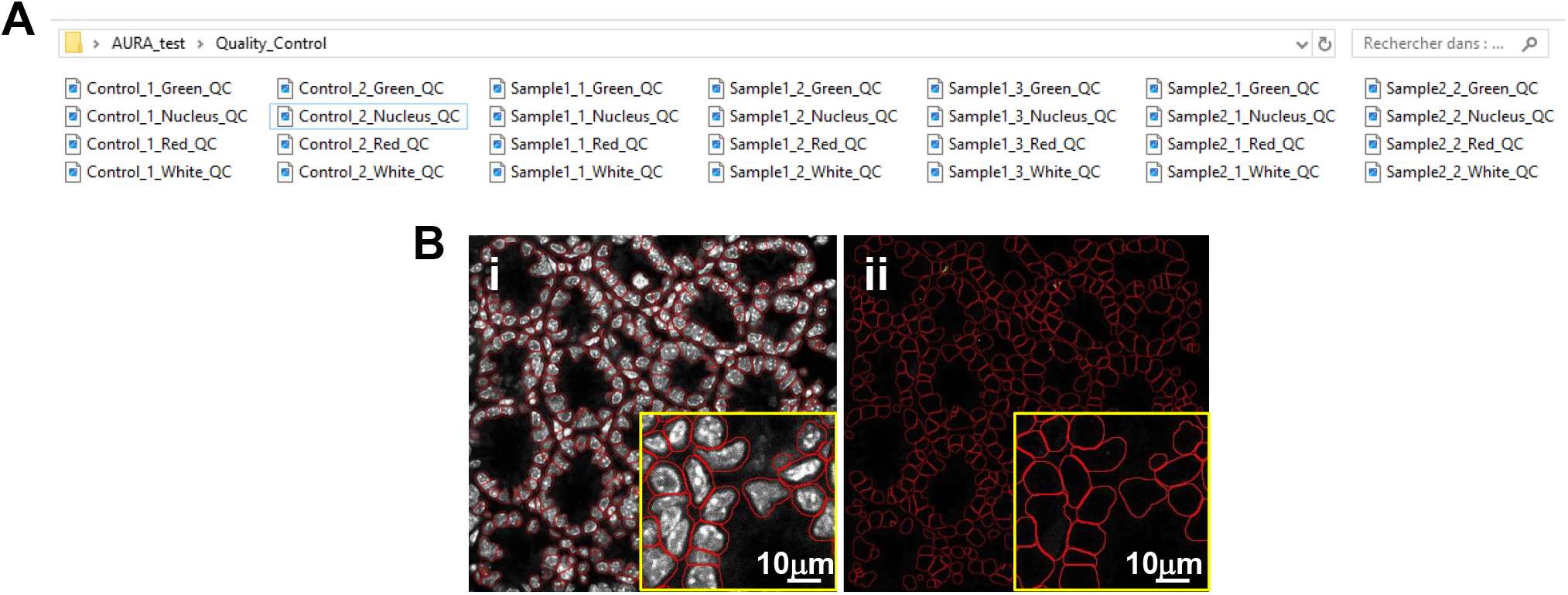
Content of the “Quality control folder”. (**A**) The subfolder contains individual quality control images for each image and analyzed channel (here Green, Nucleus, Red and White), saved as .TIFF format. Files are named according to the sample identifiers, followed by the “channel<u>_</u>name<u>_</u>QC” (e.g., “Control<u>_</u>1<u>_</u>Green<u>_</u>QC”, “Sample1_1_Nucleus_QC”). (**B**) Representative example of the quality control image showing: i) nuclei segmentation, displayed in red; (ii) Dots detection quality control. Dots are displayed in yellow with nuclei segmented in red.

**Figure 8.**
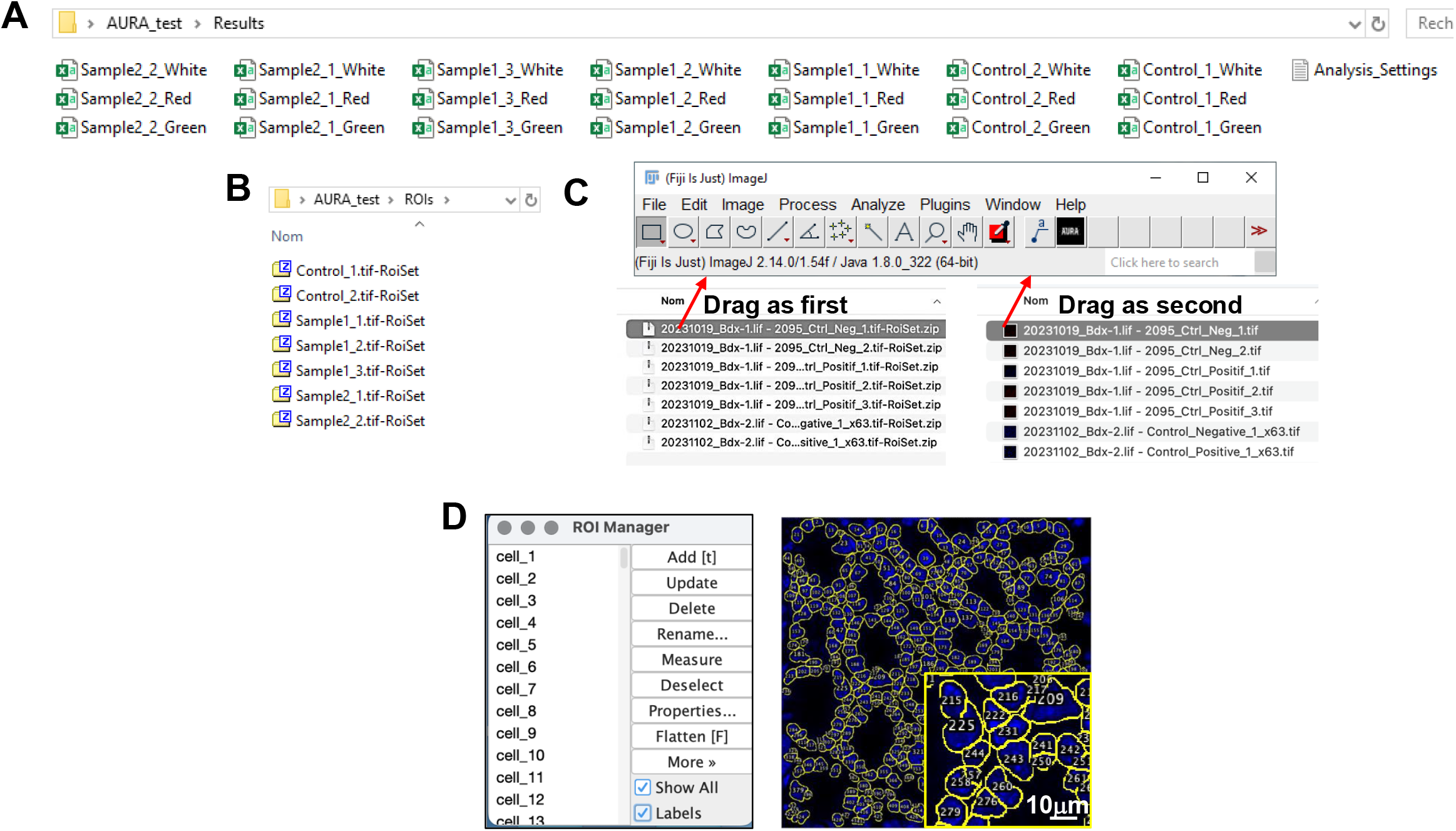
Content of the “Results” and “Region of interest” folders. **(A)** Lelt panel: the Results Folder contains the result data for each image across the different channels, saved in .CSV format. Files are named according to the image identifiers, followed by the channel name (e.g., “Sample2_1_White”, “Control_1_Red”). Right panel: the analysis settings tested are summarized in .txt file. **(B)** The ROIs Folder contains sets of regions of interest (ROIs) used in the analysis, saved as .TIFF format. Files are named according to the image identifiers (here Sample and Control), followed by “RoiSet” to indicate ROI data (e.g., “Control_1.tif-RoiSet”, “Sample1_3.tif-RoiSet”). **(C)** Open the ROI manager in Fiji. To visualize the raw data, first drag the RoiSet file, followed by the corresponding .TIFF image file for the selected channel, into the Fiji interface. **(D)** ROI manager drop-down menu (left panel). To display the segmentation, select “Show All” and “Labels” and the image with the selected ROI will open, displaying the nuclei channel (right panel). Each nucleus has an assigned number which will appear in the result file as slice # (e.g. slice 1 = Nucleus #1).

## Export & merge data

### Timing: 120 files/min

This step includes merging all .CSV files generated by the AURA macro in one single .XLSX file. This step is performed with the web app written in *Python* named “Aura Data Processing”, available at: https://aura-data-processing.streamlit.app/.

10 Set parameters for data processing
  a. Provide an experiment name without whitespace.
  b. Choose your type of analysis.
  c. Choose the input format: .CSV files or .ZIP folder.
  d. Upload your files accordingly: Drag & drop files onto the dedicated area or select files through your file manager (Browse files button). **Note:** Upload all the results .CSV files **along with the Analysis_Settings.txt** file generated by the AURA macro.
  e. Click on “Process files” to run the script. **Note**: the button “Process files” becomes clickable only after uploading your files and providing an experiment name.
  f. Follow script progression.
11 Click on “Download” to get your results in a .XLSX file. **Note:** Since Excel does not support file names that are longer than 31 characters, open the file directly with the open-source LibreOffice (https://www.libreoffice.org/) which does not have this limitation.

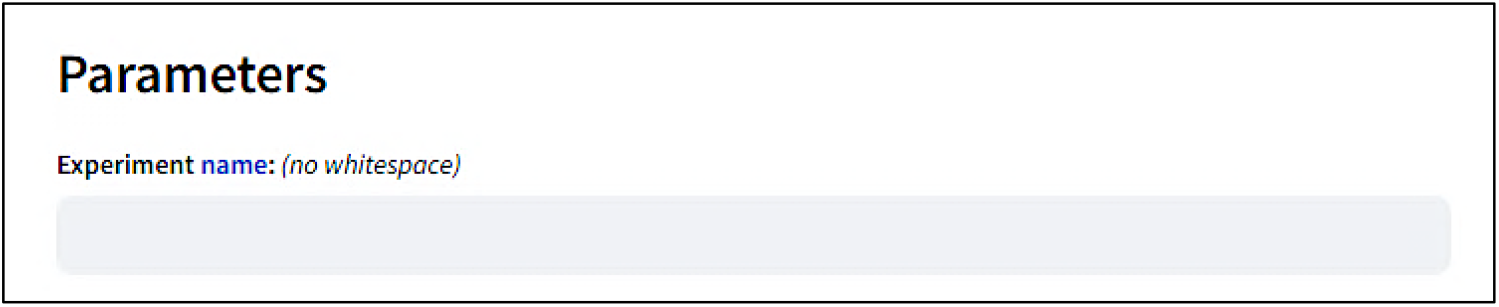

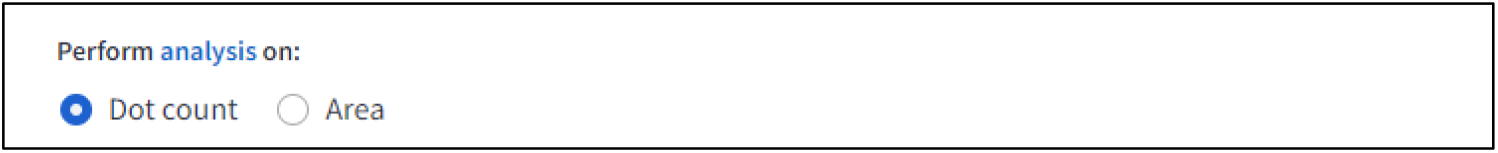

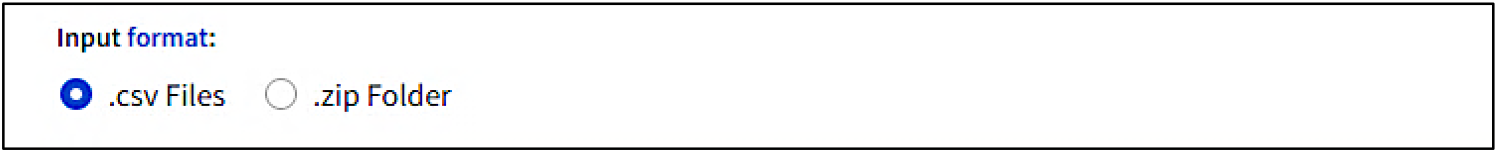

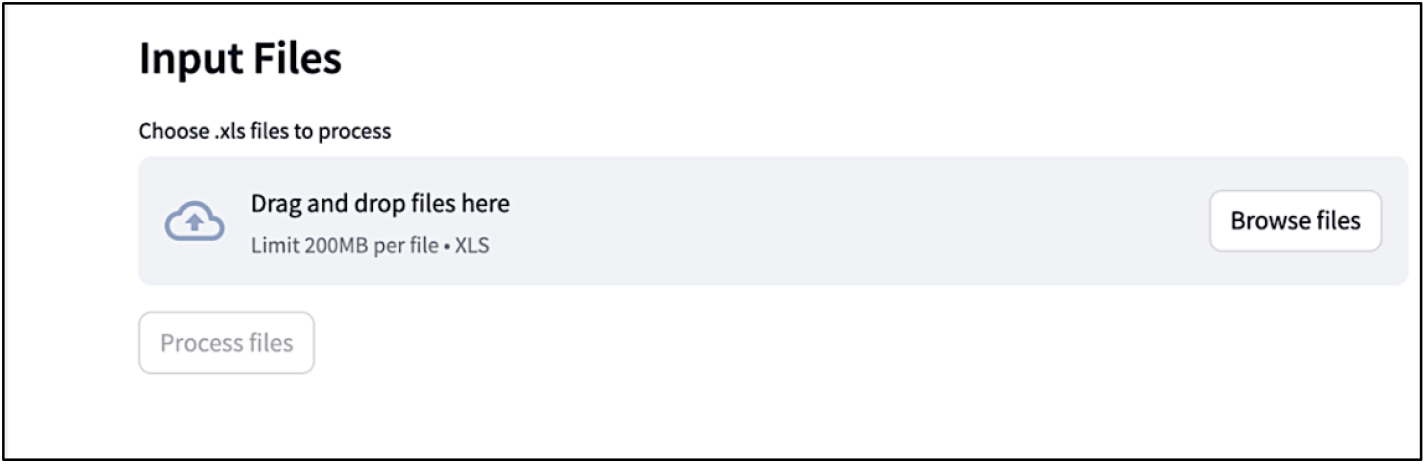

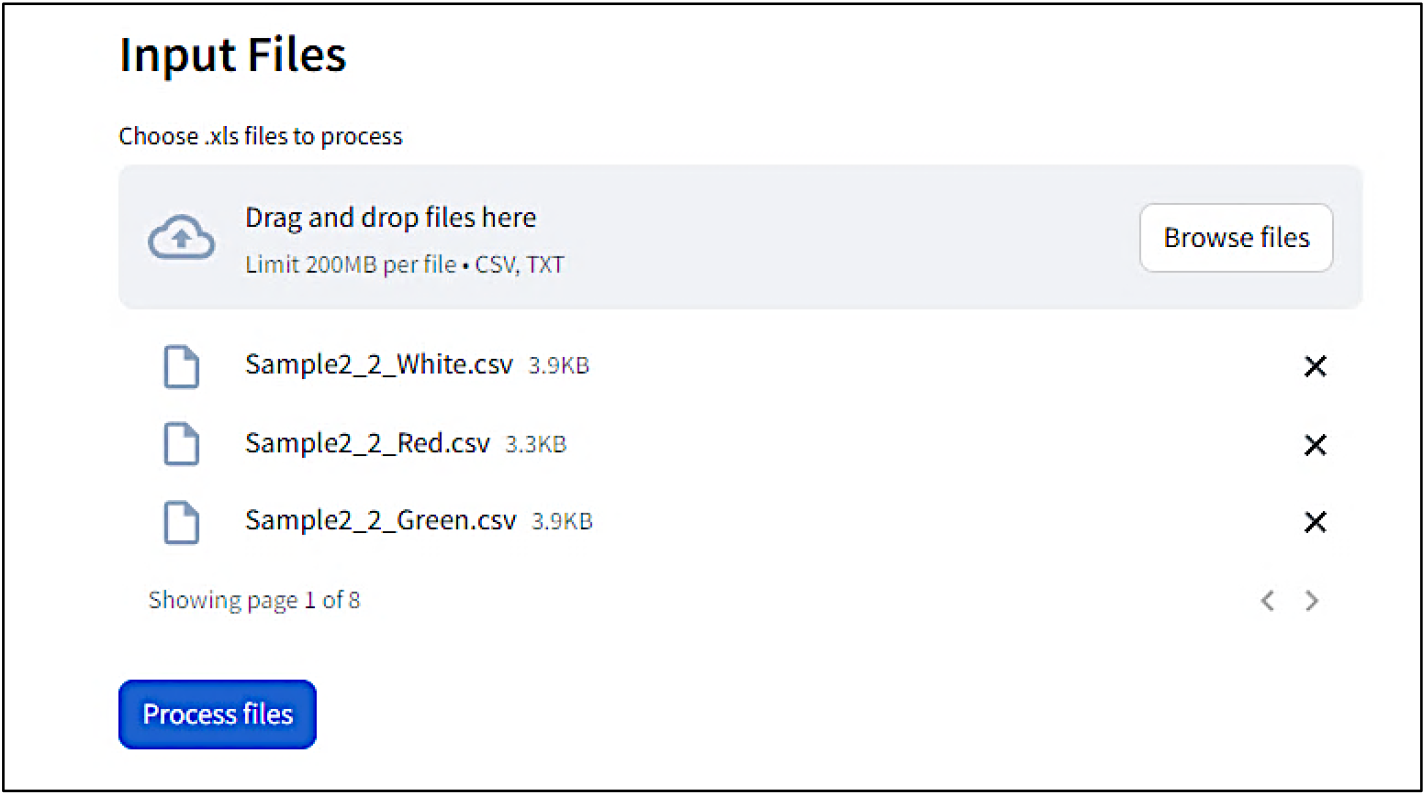

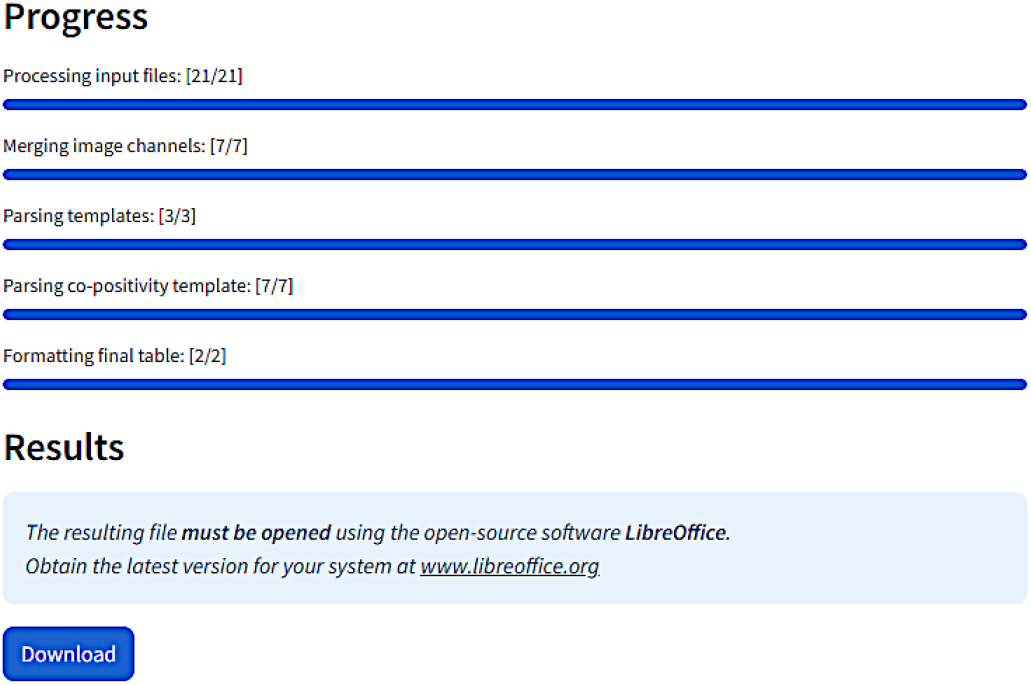

## Quality control

### Timing: 1 min by image

Use the quality control image with nuclei overlaid on the original image to inspect the segmentation quality. These images are saved in the Quality_control subfolder (step 9 and Figure 7) and should be stored for findability and accessibility ^5^.

12 Perform a visual quality control on all the analyzed images on the nuclei and dots segmentation **Note:** All nuclei segmentation images are saved as “sample_name**”_Nucleus_QC**. All dots segmentation images are saved as “sample_name”_**marker_name_QC**.

Result analysis

**Note:** Analysis of *in situ* RNA transcriptomic data can be analyzed in various ways:

a. By the percentage of positive cells.
b. By quantifying the average number of dots or transcripts per cell.
c. By co-localization based on the percentage of cells displaying one or more dots for different targets.
d. By the H-scoring ^6^. The H-score is a metric derived from RNAscope image analysis to visualize dynamic levels of a particular RNA expression within a cell population. The H-score is obtained by categorizing cells into bins based on their expression level, indicated by the number of detectable dots within a cell as follows:
  i. Bin 0 indicates no detectable staining.
  ii. Bin 1 includes cells with 1 to 3 dots.
  iii. Bin 2 comprises cells with 4 to 9 dots.
  iv. Bin 3 includes cells with 10 to 15 dots.
  v. Bin 4 represents cells with more than 15 dots.
  To measure the H-score, each bin is weighted by multiplying the percentage of cells within a bin category by the bin number. The sum of these weighted values yields the final H-score. The formula is expressed as follows:

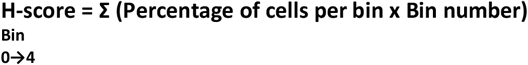
13 Open the spreadsheet containing sheets with the detailed quantification (Figure 9A) and the cumulative result (Figure 9B) for each sample. **Note:** The following information is contained in the detailed spreadsheet for each sample:
f. Number of cells per bin for each channel (Figure 9A, left panel).
  a. Percentage of positive cells per bin for each channel (Figure 9A, left panel).
  b. H-score per channel (Figure 9A, left panel).
  c. Average number of dots per cell per channel (Figure 9A, left panel).
  d. Number and percentage of double or multiple-positive cells (herein defined as copositive cells, Figure 9A, left panel).
  e. Number of dots for each cell and each channel analyzed, and co-positive cells (Figure 9A, right panel). The following information is contained in the cumulative summary spreadsheet (Figure 9B)
    a. Percentage of positive cells (Figure 9B left).
    b. H-score per channel (Figure 9B left).
    c. Average number of dots per cell per channel (Figure 9B left).
    d. Percentage of co-positive cells (Figure 9B right).

**Note**: The signal for a single mRNA molecule is detected as a punctate dot. However, multiple mRNA molecules localized in close proximity can result in a cluster. This cluster is handled indirectly. In step 10b, choose Area as Analysis type, and the result file will contain the percentage of positive cells and the area occupied by the probe signal per cell instead of the number of dots (Figure 10).

**Note:** The protocol has been validated by re-analyzing published manually curated data sets from the Frisan’s laboratory ^7^. The mRNA levels for the two inflammatory sensors AIM2 and NLRP3 were investigated in the colon and liver of mice either left uninfected or infected with the *Salmonella enterica*, serovar Typhimurium strain MC1 expressing a functional (MC1 TT) or non-active (MC1 Δ*cdtB*) typhoid toxin. The manual and AURA-curated results are comparable, showing similar statistical differences among the samples, which supports the validity of the application. However, manual curation took approximately 3.5 weeks, while the AURA evaluation was completed in approximately 60 minutes (Figure 11). Additionally, manual curation requires merging the different channels before starting the counting.

**Note:** AURA can potentially be applied to analyze chromogenic images of *in situ* mRNA hybridization. This step requires color separation to separate the markers into different color channels in Fiji, which is done as follows:

a. Images need to be in the RGB format: in the Fiji menu navigate to Image > Type and scroll down to select RGB color.
b. In the Fiji menu navigate to Image > Color and scroll down to select Color Deconvolution.
c. From the Color Deconvolution dropdown menu select “From ROI” (scroll down if it is not visible among the first choices).
d. Use the rectangle tool to select examples of nuclei, probes, and background stains.
e. Run the color deconvolution to generate separate images for each channel. Save images in .GIF format in a folder and proceed with the AURA protocol.

**Figure 9.**
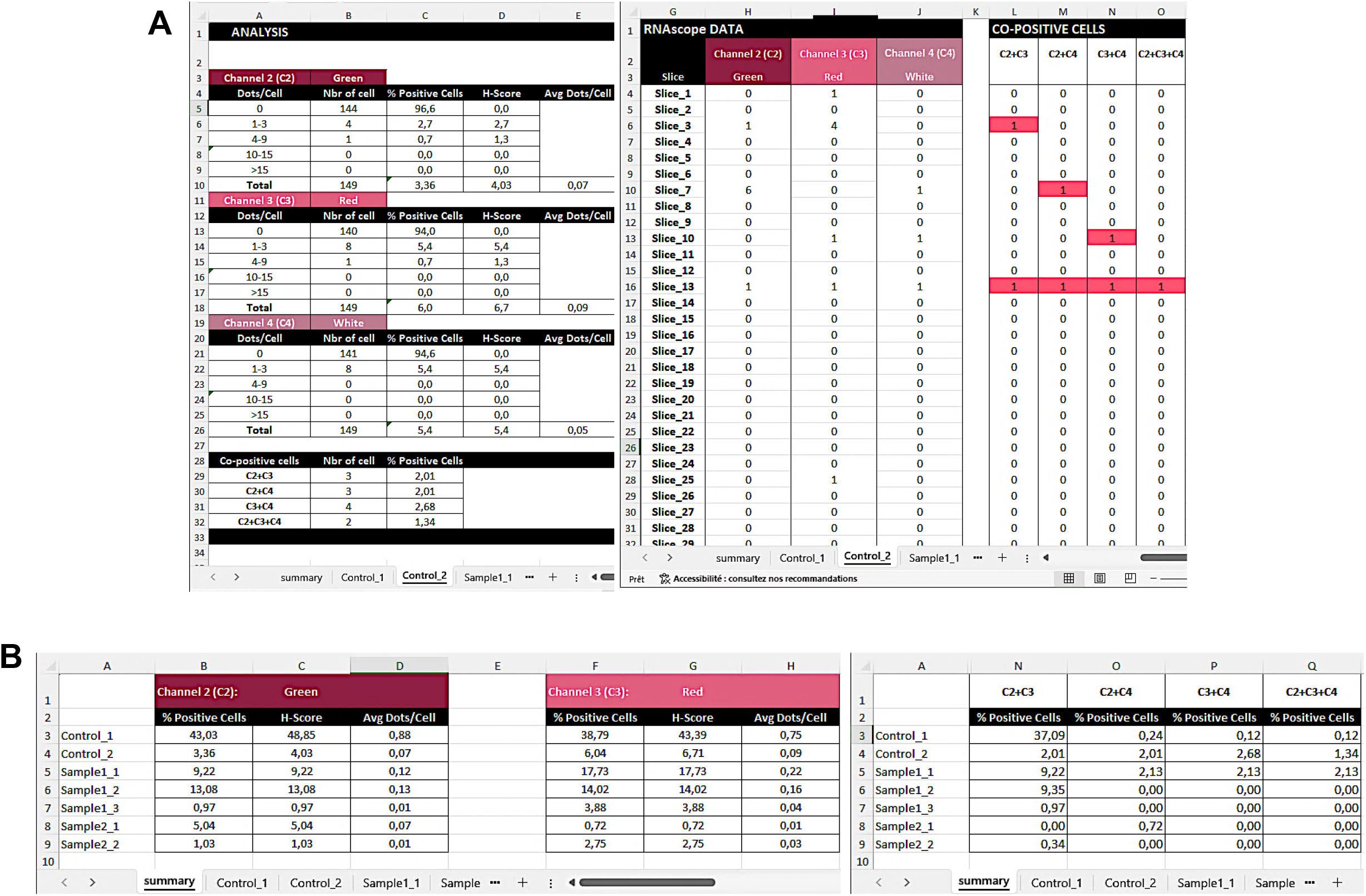
Example output summary data files. (**A**) Summary for each sample. Left panel: representative spreadsheet reporting the number of cells in each of the bin categories, the percentage of positive cells per bin category, the H-score, and the average number of dots per cell (Avg Dots/Cell) per each acquired channel (here Green, Red, and White), as well as the number and percentage of co-positive cells in one selected sample (Control_2). Right panel: spreadsheet containing information for the number of dots per nucleus in each channel (slice 1 = nucleus 1) and the number of dots per nucleus for co-positive cells in the three analyzed channels in one selected sample (Control<u>_</u>2). Co-positive cells are highlighted in colour. (**B**) Consolidate summary for all samples in a spreadsheet presenting the percentage of positive cells, the H-score, the average number of dots per cell for each sample analysed in the single channels, and the percentage co-positive cells in the three channels analyzed.

**Figure 10.**
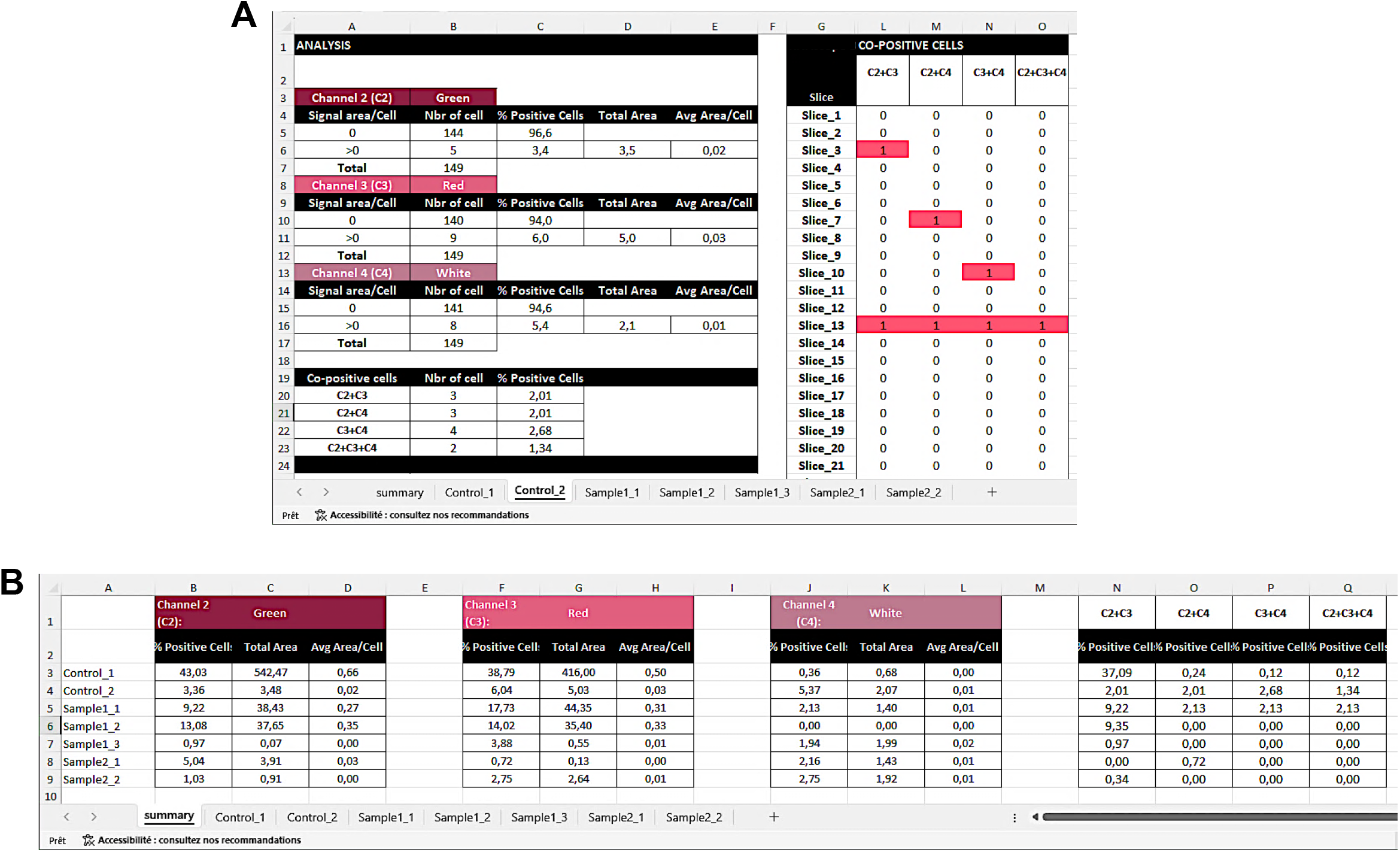
Example output summary data files using the area as analysis parameter. (**A**) Summary for each sample. Left panel. Representative spreadsheet reporting the number of cells counted, the percentage of positive cells per each acquired channel (here Green, Red, and White), total area covered by the dot/cluster fluorescence, the average of dot/cluster fluorescent area per cell (Avg Area/cell), as well as the number and percentage of co-positive cells in one selected sample (Control_2). Right panel: representative spreadsheet containing information on the co-positive cells (slice 1 = nucleus 1) in one selected sample (Control_2). Co-positive cells are highlighted in colour. **(B)** Consolidate summary for all samples in a spreadsheet presenting the percentage of positive, the total area of the dot/cluster fluorescent area, the average of dot/cluster fluorescent area per cell in the single channels, and the percentage co-positive cells in the three channels analyzed.

**Figure 11.**
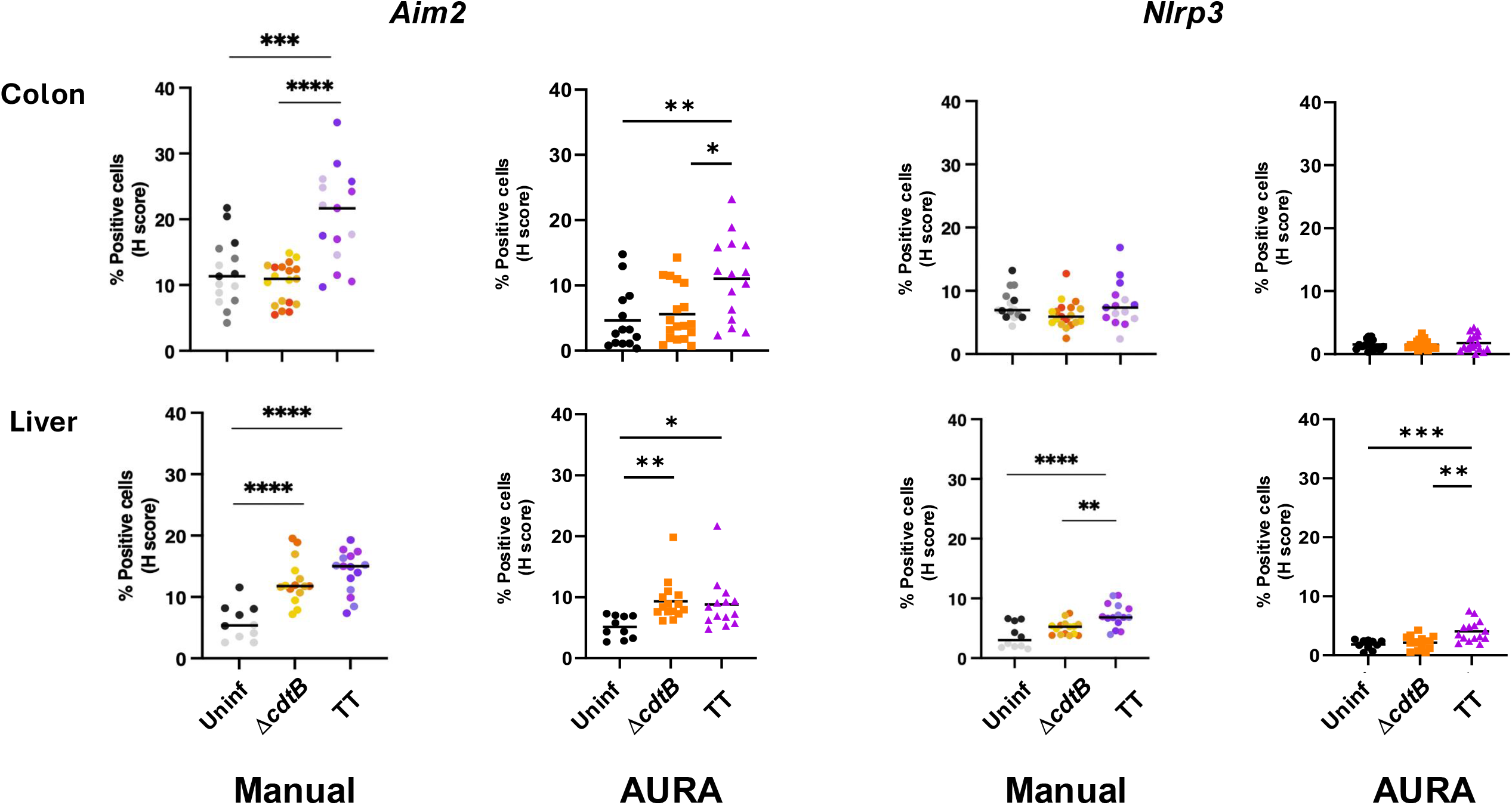
Comparison of manual versus AURA quantification. Mice were mock-treated with PBS (Uninf) or infected with *Salmonella* MC1 Δ*cdtB* (Δ*cdtB*) or MC1 TT (TT) for 10 days. The mRNA levels of the inflammasome sensors AIM2 and NLRP3 were assessed in the colon and liver by RNAscope™ and manually curated (manual) or analyzed with AURA (AURA). The data are presented as percentage positive cells in all the micrographs acquired for each mouse. * p-value ≤ 0.05; ** *p*-value ≤ 0.01; *** *p*-value ≤ 0.001; **** *p*-value ≤ 0.0001.

### Expected outcomes

This protocol provides a straightforward, open-source, cost-free, and user-friendly macro/script designed to analyze large volumes of *in situ* RNA hybridization microscopy data. The output includes segmented nuclei and dot identification, which are saved as separate files. The script combines separated .CSV files into a single .XLSX file, with a pre-analysis for each sample and a final summary analysis for all samples. Quality control images are saved to visually validate segmentation accuracy. Additionally, this method allows for cell-by-cell control and facilitates data storage and management in line with findability and accessibility guidelines ^5^.

### Quantification and statistical analysis

The quantification of the percentage of single or multiple positive cells is directly provided in the result output spreadsheet (step 13 of the “Step-by-Step method details” section). Statistical analyses can be performed using Prism 7 and Graph Pad Software, or any statistical analysis software, applying the appropriate statistical methods depending on the experimental set-up, e.g. comparing two or multiple experimental groups.

### Limitations

Nuclei staining and imaging quality

Nuclei staining and image acquisition quality are crucial for accurate analysis. Good nuclei staining and image acquisition should exhibit low background noise and minimal overlap between nuclei. Poor-quality staining and image capturing can lead to incomplete segmentation and unreliable quantification. Figure 12 provides examples of poor and good-quality nucleus staining and imaging, with the corresponding nuclei segmentation results.

**Figure 12.**
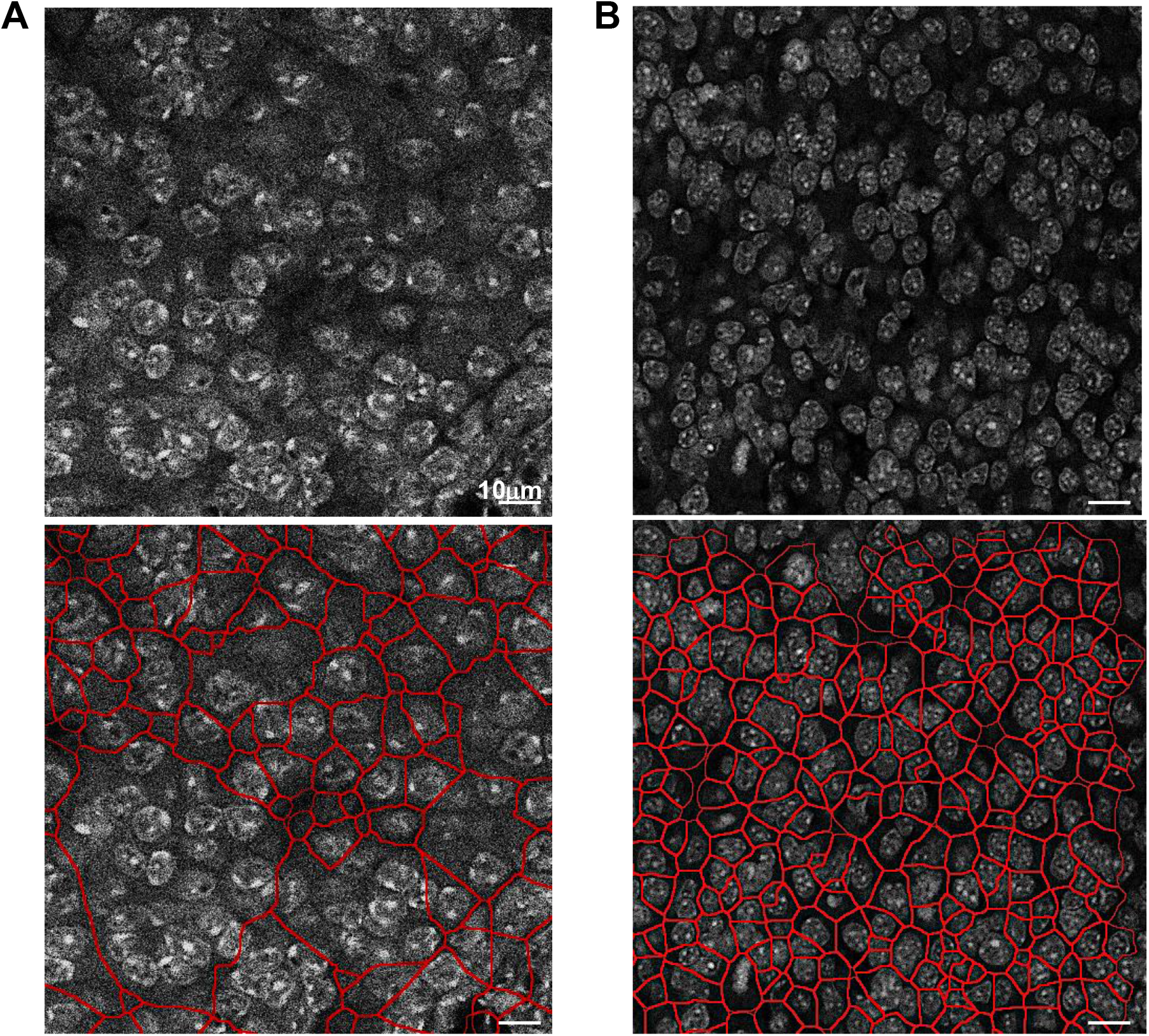
Examples of nucleus staining image files. Representative micrographs of DAPI stained nuclei of bad quality **(A)** where the nuclei are not well defined, and the image has a high background/signal ratio, and good quality image with a very defined nuclei staining and low background /signal ratio **(B)**. The lower panel shows the results of the AURA nuclei segmentation.

### Troubleshooting

#### Problem 1

Error message when starting the AURA Light or AURA macro (related to steps 2 and 6 of the “Step-by-step method details” section).

This may occur if all the plugins are not installed or if special characters were used when naming your channels.

#### Potential solution

- Ensure that all the plugins (downloaded in step 6c “Before you begin” section) are installed.
- Avoid using special characters for the channels’ names.

#### Problem 2

Using a zero value for nuclei expansion results in poor segmentation (related to steps 4d and 7d of the “Step-by-step method details” section).

#### Potential solution

The macro considers a value of 0 as infinite expansion; hence the default value is set at 0,1µm.

#### Problem 3

Inaccurate nuclei segmentation with AURA (related to step 7 of the “Step-by-step method details” section).

#### Potential solution

Before you start the batch analysis we recommend using AURA Light (steps 1 to 5 “Step-by-step method details” section), testing several parameters for nuclei and dot segmentation, and cell expansion. If the segmentation is still not satisfactory, we propose a list of parameters that can be modified in the AURA and AURA Light macro code (Table 1 and Table 2). The code lines are the same for both macros.

**Table 1:** Key parameters for nuclear segmentation in AURA light and AURA macro code.

| Nuclei segmentation Parameters | Code modifications examples | AURA Line |
| --- | --- | --- |
| Increase saturation | run("Enhance Contrast", "saturated=0.50"); | 27 |
| Decrease radius | run("Mean...", "radius=1.5"); | 29 |
| Increase gaussian blur | run("Gaussian Blur...", "sigma=3"); | 30 |
| Add image sharpening | run("Sharpen"); | 31 |
| Use other automatic threshold (Otsu, Huang, Yen, etc) | setAutoThreshold("Huang dark"); | 31 |
| Changes in post-processing steps | run("Dilate"); run("Invert"); run("Close"); | 33 |
| Add despeckle after watershade | run("Despeckle"); | 35 |
| Apply Watershed before Mask conversion | run("Watershed"); | 32 |

**Table 2:** Key parameters for dot segmentation in AURA light and AURA macro code.

| **Dot segmentation parameters** | **Code modifications examples** | **AURA Line** |
| --- | --- | --- |
| Increase saturation | `run("Enhance Contrast", "saturated=0.60");` | 43 |
| Increase gaussian blur | `run("Gaussian Blur...", "sigma=1");` | 44 |
| Replace gaussian blurr with a median filter | `run("Median...", "radius=2");` | 44 |
| Apply Watershed before Mask conversion | `run("Watershed");` | 46 |
| Changes in post-processing steps | `run("Dilate"); run("Close"); run("Invert");` | 47-49 |

Figure 13 illustrates the changes in the nuclear segmentation when the parameters highlighted in bold in the figure have been modified. The data show that, regardless of the code used, segmentation errors are present, which is an intrinsic limitation of batch analysis. This problem can be mitigated by acquiring a large number of nuclei.

**Figure 13.**
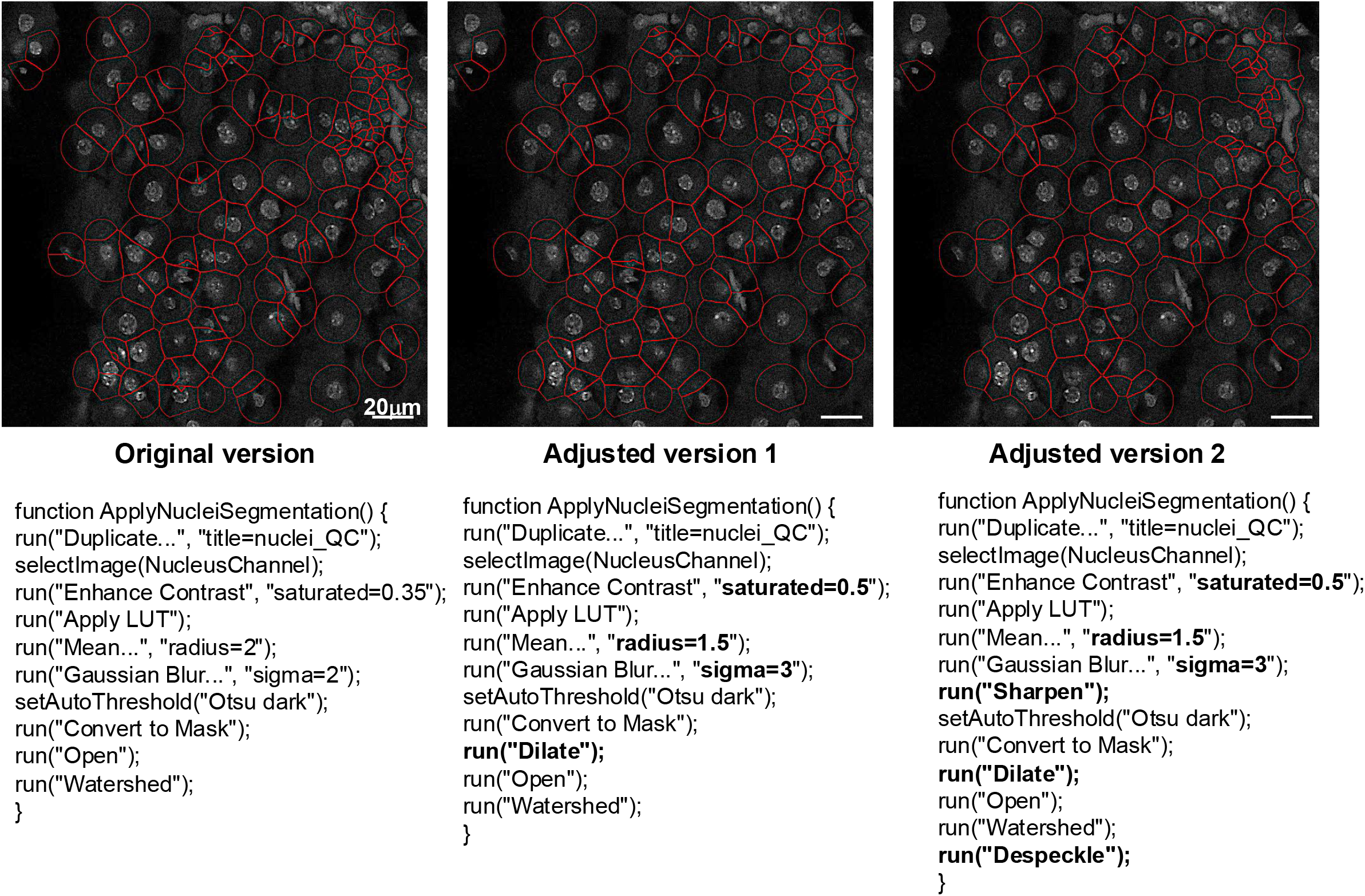
Nucleus segmentation results upon changing the code parameters. Representative micrographs of DAPI stained nuclei (upper panel) where the code modifications (lower panel) highlighted in bold have been applied.

In addition, we provide below critical code lines for the nuclei and dots segmentation. This step requires Fiji coding expertise.

- Parameters for nucleus segmentation
- Parameters for defining cell area
- Parameters for dots segmentation
- Parameter for dot analysis. This step is repeated automatically for each channel you identified in Step 4 of the “Step-by-step method details” section.

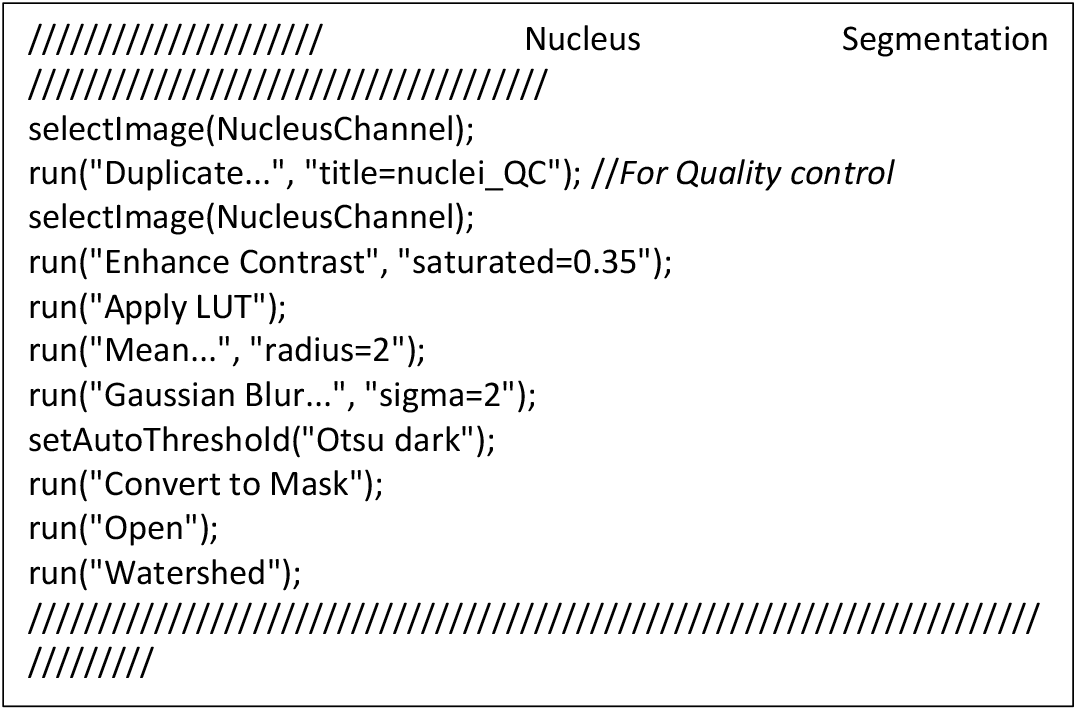

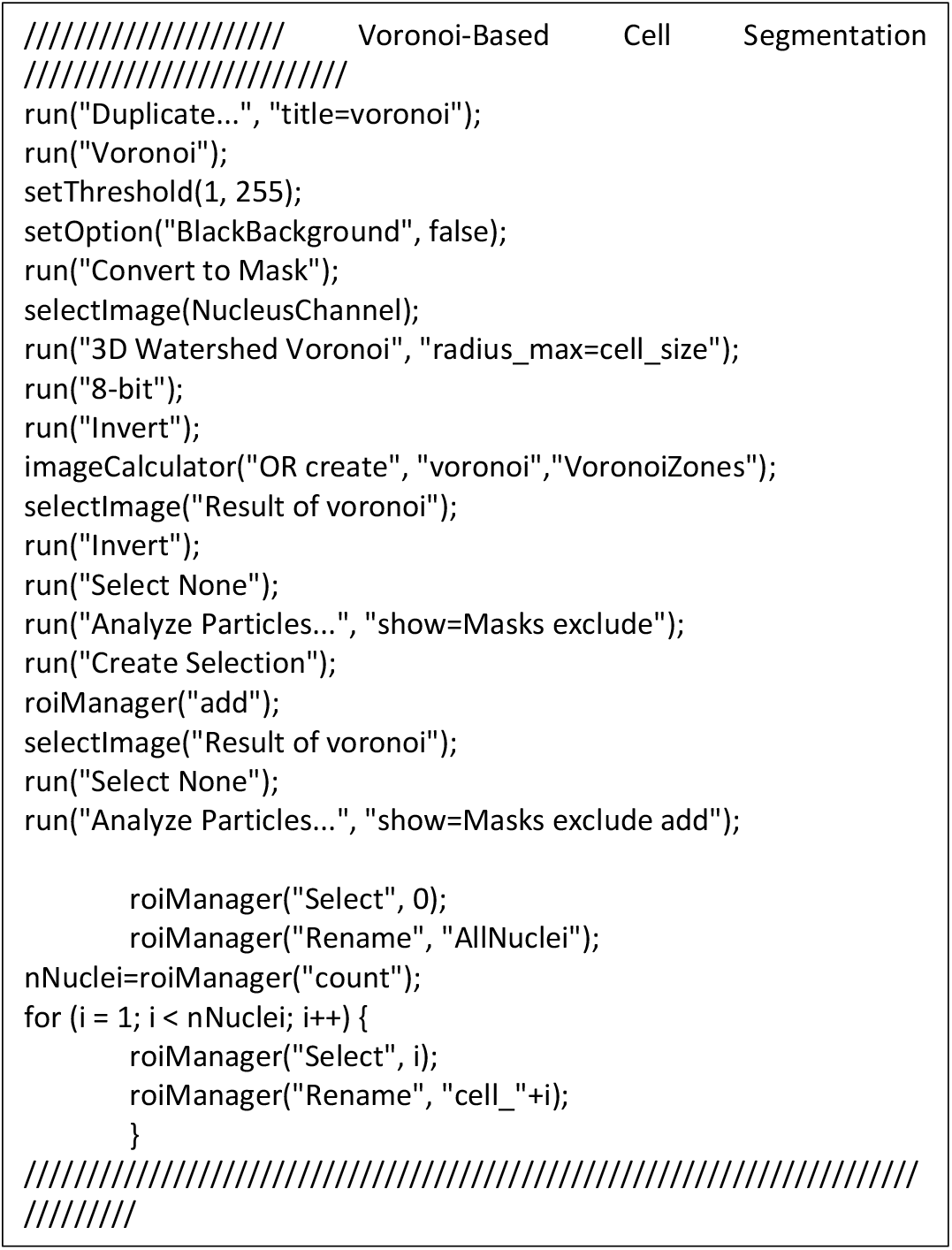

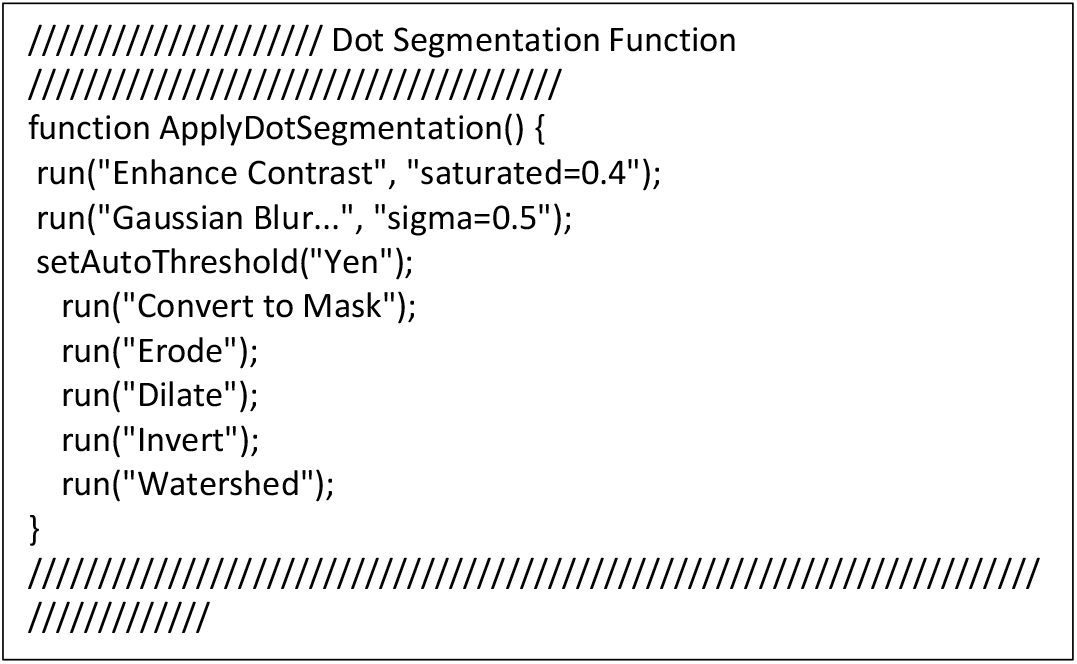

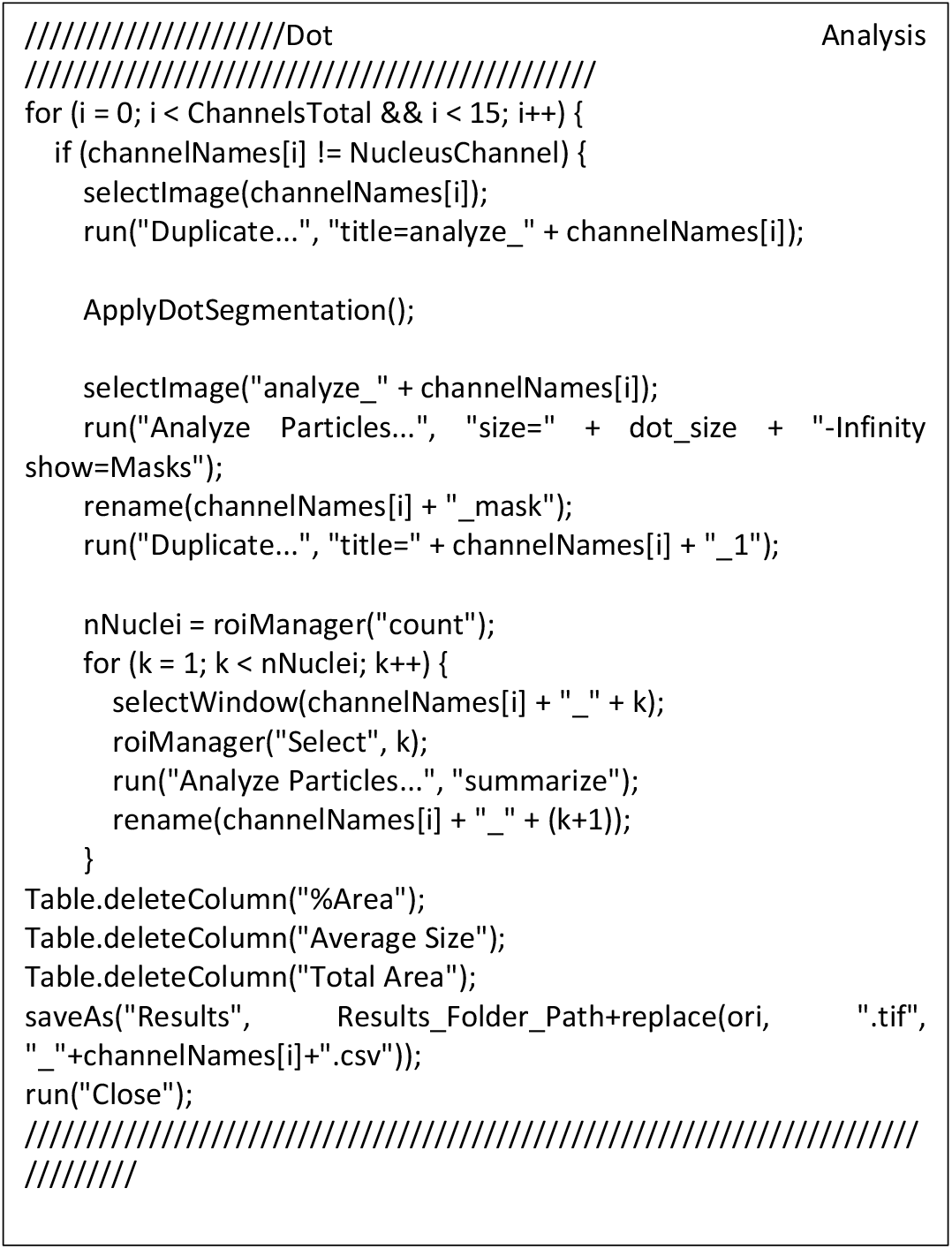

If inaccurate segmentation is the result of poor nucleus staining or image quality, you can use the following steps:

- Make sure to exclude any images from your analysis if they are unusable. It is recommended to take multiple images to ensure a sufficient number of high-quality images for the analysis (700 to 1000 high-quality nuclei per sample).
- Perform empirical analysis by manually counting the nuclei while retaining the dots segmentation.
- Retake images of your sample, ensuring better nuclei staining and nuclei acquisition.

#### Problem 4

Presence of clusters (related to step 7f of the “Step-by-step method details” section).

#### Potential solution

Abundant transcripts could generate clusters of “dots”. The cluster is handled indirectly in step 10b of the “Step-by-step method details” section where Area is chosen as the Analysis type (Note in step 10b).

In case mRNA targets show different patterns (low expression with distinguishable dots and high expression with clusters), we recommend running the AURA macro as described in steps 6 to 8 of the “Step-by-step method details” section for all probes. Then, perform a separate analysis using the *Python* application as follows:

- Upload the .CSV results for the low-expressed probe and the Setting_Analysis.txt file, and run the analysis choosing the “dot” mode (Step 10b of the “Step-by-step method details” section).
- Upload the .CSV results for the highly expressed target and the Setting_Analysis.txt file, and run the analysis choosing the “area” mode (Step 10b of the “Step-by-step method details” section).
- Merge the data from the two summary spreadsheet files for the samples of interest. This procedure will limit the information about the H-score for one sample set, but it will allow the user to retrieve information regarding the percentage of positive and co-positive cells.

#### Problem 5

Dots are not counted by the AURA macro (related to steps 4f and 7f of the “Step-by-step method details” section). This can happen if the threshold set for dot detection is too high.

#### Potential solution

- Reduce the threshold for dot detection to ensure more dots are counted (steps 4f and 7f of the “Step-by-step method details” section).
- Adjust the relevant macro code at lines 40-51 to fine-tune dot detection parameters (Table 2).

#### Problem 6

Remembering analysis parameters (related to steps 4 and 7 of the “Step-by-step method details” section). Failure to recall parameters may render the process of replicating or reviewing the analysis more complex.

#### Potential solution

Refer to the file named Setting_Analysis.txt in the Results folder, where settings parameters are stored (Figure 4A).

#### Problem 7

Error message when connecting to AURA Data Processing *Python* App (related to step 10 of the “Step-by-step method details” section).

#### Potential solution

The *Python* App will automatically enter sleep mode after several days of inactivity. Click the refresh button and wait a few minutes for the app to start.

#### Problem 8

AURA Data Processing *Python* App does not process files (related to step 10 of the “Step-by-step method details” section).

#### Potential solution

- Ensure that you have uploaded the Setting_Analysis.txt file, which distinguishes files with similar names.
- Try uploading your results folders as a .ZIP file to ensure that all necessary files are included.

#### Problem 9

Issues on the summary page when opened with Microsoft Excel (related to step 11 of the “Step-by-step method details” section). The problem occurs if your data file names are longer than 31 characters, which is not supported in Microsoft Excel.

#### Potential solution

Open your file with LibreOffice Calc, which does not have this limitation.

## Resource availability

### Lead contact

Further information and requests for resources should be directed to and will be fulfilled by the lead contact, Jean Descarpentrie

### Technical contact

Technical questions on executing this protocol should be directed to and will be answered by the technical contact, Wilfried Souleyreau.

### Material availability

This kind of study did not generate unique products.

### Data and code availability

The AURA Light and AURA macro are available at https://github.com/FlorianBrnrd/aura-data-processing.

The web app “AURA data processing”, which allows users to merge .CSV files into a unique

.XLSX file, can be found at https://aura-data-processing.streamlit.app/.

Codes are available at https://github.com/FlorianBrnrd/aura-data-processing.

The GitHub repo has been archived at Zenodo: https://zenodo.org/records/13908524. Zenodo DOI: 10.5281/zenodo.13908524

## Acknowledgments

This investigation was supported by grants from The Swedish Research Council (2021-00960), The Swedish cancer Society (23 2814 Pj) the Kempestiftelserna (2021 JCK-3110), the Cancer Research Foundation in Northern Sweden (AMP20-993), and Umeå University (to T.F.). We acknowledge the Biochemical Imaging Center Umeå (BICU) at Umea University and theNational Microscopy Infrastructure (NMI) (VR-RFI 2016-00968) for assisting with microscopy.

## Author contributions

Conceptualization, O.C.B.M., W.S.; methodology, J.D., W.S.; app design, F.B.; data production MLC, writing, J.D., W.S., TF; funding acquisition and supervision, LB, TM, I.S.P., T.F.

## Declaration of interests

The authors declare no competing interests.

## Notes

### Competing Interest Statement

The authors have declared no competing interest.

### Summary of Updates

A more detailed protocol description has been added in response to the reviewers comments. The manuscript has been accepted in principle by STAR Protocols.

## References

1. Patel, A., Balis, U.G.J., Cheng, J., Li, Z., Lujan, G., McClintock, D.S., Pantanowitz, L., and Parwani, A. (2021). Contemporary Whole Slide Imaging Devices and Their Applications within the Modern Pathology Department: A Selected Hardware Review. J Pathol Inform 12, 50. 10.4103/jpi.jpi_66_21.

2. Schindelin, J., Arganda-Carreras, I., Frise, E., Kaynig, V., Longair, M., Pietzsch, T., Preibisch, S., Rueden, C., Saalfeld, S., Schmid, B., et al. (2012). Fiji: an open-source platform for biological-image analysis. Nature methods 9, 676–682. 10.1038/nmeth.2019.

3. Schneider, C.A., Rasband, W.S., and Eliceiri, K.W. (2012). NIH Image to ImageJ: 25 years of image analysis. Nature methods 9, 671–675. 10.1038/nmeth.2089.

4. Ollion, J., Cochennec, J., Loll, F., Escude, C., and Boudier, T. (2013). TANGO: a generic tool for high-throughput 3D image analysis for studying nuclear organization. Bioinformatics 29, 1840–1841. 10.1093/bioinformatics/btt276.

5. Wilkinson, M.D., Dumontier, M., Aalbersberg, I.J., Appleton, G., Axton, M., Baak, A., Blomberg, N., Boiten, J.W., Santos, L.B.D., Bourne, P.E., et al. (2019). The FAIR Guiding Principles for scientific data management and stewardship (vol 15, 160018, 2016). Sci Data 6. ARTN 6 10.1038/s41597-019-0009-6.

6. Gaide, N., Crispo, M., Jbenyeni, A., Bleuart, C., Delverdier, M., Vergne, T., Le Loc’h, G., and Guerin, J.L. (2023). Validation of an RNAscope assay for the detection of avian influenza A virus. J Vet Diagn Invest 35, 500–506. 10.1177/10406387231182385.

7. Chiloeches, M.L., Bergonzini, A., Martin, O.C.B., Bergstein, N., Erttmann, S.F., Aung, K.M., Gekara, N.O., Cariño, J.F.A., Pateras, I.S., and Frisan, T. (2024). Genotoxin-producing induces tissue-specific types of DNA damage and DNA damage response outcomes. Frontiers in immunology 14. ARTN 1270449 10.3389/fimmu.2023.1270449.

